# Metagenomic analysis of the cow, sheep, reindeer and red deer rumen

**DOI:** 10.1101/2020.02.12.945139

**Authors:** Laura Glendinning, Buğra Genç, R John Wallace, Mick Watson

**Affiliations:** Genetics and Genomics, The Roslin Institute and Royal (Dick) School of Veterinary Studies, University of Edinburgh, Edinburgh, Midlothian, United Kingdom; Rowett Institute, University of Aberdeen, Ashgrove Rd W, Aberdeen, Aberdeenshire, United Kingdom

**Keywords:** microbiota, deer, cow, sheep, reindeer, mag

## Abstract

The rumen microbiota comprises a community of microorganisms which specialise in the degradation of complex carbohydrates from plant-based feed. These microbes play a highly important role in ruminant nutrition and could also act as sources of industrially useful enzymes. In this study, we performed a metagenomic analysis of samples taken from the ruminal contents of cattle (*Bos Taurus*), sheep (*Ovis aries*), reindeer (*Rangifer tarandus*) and red deer (*Cervus elaphus*). We constructed 391 metagenome-assembled genomes originating from 16 microbial phyla. We compared our genomes to other publically available microbial genomes and found that they contained 279 novel species. We also found significant differences between the microbiota of different ruminant species in terms of the abundance of microbial taxonomies, carbohydrate-active enzyme genes and KEGG orthologs. However, we found that the vast majority of carbohydrate-active enzymes were present in all of our sample types, which may indicate that there is a core set of these enzymes which are present across ruminants and are independent of diet and environmental conditions. We present a dataset of rumen-derived genomes which in combination with other publicly-available rumen genomes can be used as a reference dataset in future metagenomic studies.

**Data Summary:** The paired-read fastq files supporting the conclusions of this article are available in the European Nucleotide Archive repository (https://www.ebi.ac.uk/ena/browser/view/PRJEB34458). The RUG fasta files supporting the conclusions of this article are available in the Edinburgh DataShare repository (https://doi.org/10.7488/ds/2640).

## Introduction

The microbial communities which inhabit the rumen contain a mixture of bacteria, fungi, protozoa, viruses and archaea, and through fermentation are able to convert complex plant carbohydrates into short-chain volatile fatty acids. The microbial pathways present have a large impact on feed efficiency (1–3), alongside other important production traits such as milk and fat yield (4, 5). Understanding the processes by which food is digested in the rumen may allow us to improve feed efficiency in ruminants (1), either by the production of enzymes isolated from microbes (6) or by manipulating the microbiota through the use of pre- or probiotics (7). There are also other potential industrial uses for the proteins produced by ruminal microbes, for example in processing biofuels, bioremediation, processing pulp/paper and textile manufacturing (8–11). Ruminants are also the largest source of anthropogenic methane emissions and gaining a greater understanding of which microbes are important in methane production could lead to improved methane mitigation strategies (7,12–16).

While inroads have been made towards culturing members of the ruminal microbiota (17, 18) there are still many members which have not been characterised. Metagenomics is a powerful tool which allows us to examine the entire genetic repertoire of the rumen microbiota without the need for culturing. Our group has previously published thousands of metagenome assembled genomes from cattle rumen samples (19, 20) and hundreds of genomes from chicken caecal samples (21), many of which were identified as novel species.

Several studies have examined the rumen microbiota using metagenomic techniques in cattle and sheep; however, less effort has been made to characterise the microbiota of other ruminant species which may be less commercially-important but which could harbour microbes which could be industrially useful. For example, wild ruminants are likely to consume a far more diverse diet than farm-raised individuals, and are therefore likely to contain microbes which are able to digest different substrates. In this paper we analyse rumen metagenomic data from four ruminant species: cattle (*Bos Taurus*), sheep (*Ovis aries*), red deer (*Cervus elaphus*) and reindeer (*Rangifer tarandus*). We compare the microbiota of these species taxonomically and functionally and construct 391 named rumen-uncultured genomes (RUGs), representing 372 putative novel strains and 279 putative novel species.

## Methods

### Experimental design

Reindeer (*Rangifer tarandus*: Grazing mixed vegetation, n=2) and red deer (*Cervus elaphus*: Grazing mixed vegetation, n=4) were shot in the wild, and ruminal digesta samples were collected immediately. Samples were taken from Holstein cattle (*Bos Taurus*: Fed total mixed ration (once a day), n=4) and Finn-Dorset cross sheep (*Ovis aries*: Grazing mixed pasture, n=2) via a rumen cannula. Samples were taken from sheep after morning grazing. Sheep sampling was performed as described in McKain et al. (22). Cattle samples were taken 3 hours post feeding. Samples were collected from the bovine rumen in the following locations: top near cannula, middle at the front of the rumen, middle towards the back of the rumen and bottom (approximately 45cm down from the entrance to the rumen). Digesta samples were mixed with buffer containing glycerol as a cryoprotectant (22). The mixtures were kept on ice for 1-2 hours then frozen at −20°C. DNA extraction was performed using repeated bead beating plus column filtration, as described in (23). Shotgun sequencing was performed on an Ilumina Hiseq 2000, producing an average of 1626 million paired reads per sample, of 100bp or 150bp in length.

### Bioinformatics

Illumina adaptors were removed using trimmomatic (24) (v.0.36). IDBA-UD (25) (v.1.1.3) with the options --num_threads 16 --pre_correction --min_contig 300 was used to perform single sample assemblies. After indexing using BWA index (v.0.7.15), BWA-MEM was used to map reads to assemblies (26). BAM files were created by SAM tools (27) (v.1.3.1) and coverage was calculated using the command jgi_summarize_bam_contig_depths from the MetaBAT2 (v.2.11.1) software package (28). A coassembly was carried out on all samples using MEGAHIT (29) (v.1.1.1) with the options --continue --kmin-1pass -m 100e+10 --k-list 27,37,47,57,67,77,87 --min-contig-len 1000 -t 16. After filtering out reads which were <2kb, indexing and mapping were performed as for single assemblies.

Metagenomic binning was carried out using MetaBAT2 with the options --minContigLength 2000, -- minContigDepth 2. From the single-assemblies, 1691 bins were created and from the co-assembly 2508 bins were created. Completeness and contamination of bins were calculated using CheckM (options: lineage_wf, -t 16, -x fa) (v.1.0.5), and the bins were dereplicated using dRep (30) (options: dereplicate_wf -p 16 -comp 80 -con 10 -str 100 –strW 0) (v.1.1.2). Thus, bins were discarded if their completeness was <80% or if they had contamination >10%. The dereplicated ‘winning’ bins are referred to below as RUGs. MAGpy was used to compare the RUGs to public datasets (31).

Taxonomies were assigned to MAGs using GTDB-Tk (32). Trees produced by MAGpy were rerooted at the branch between archaea and bacteria using Figtree (33) (v.1.4.4) and visualised using GraPhlAn (34) (v.0.9.7). For submission to public repositories, our RUGs were named as the lowest taxonomic level at which NCBI and GTDB-Tk matched. The taxonomies assigned to RUGs were manually checked against the taxonomic tree and improved accordingly.

Carbohydrate-active enzymes (CAZymes) were identified using dbCAN2 (version 7, 24^th^ August 2018) by comparing RUG proteins to the CAZy database (35). RUG proteins were compared to the KEGG database (downloaded on Sept 15th 2018) (36) using DIAMOND (37) (v0.9.21). KEGG hits for which the alignment length was ≥90% of the query length were retained. The likely KEGG ortholog group for each RUG protein was inferred from the DIAMOND search results and the KEGG database. CAZyme and KEGG ortholog abundances were calculated as the sum of the reads mapping to RUG proteins within each group after using DIAMOND to align reads to the RUG proteins. PULpy was used to identify polysaccharide utilisation loci (38).

Statistical analyses were carried out within R (version 3.5.1). The ggplot2 (39) package was used to construct scatter plots and NMDS graphs. The vegan package (40) was used to create NMDS axes using the Bray–Curtis dissimilarity. The Adonis function from the vegan package was used to perform PERMANOVA analyses and DeSeq2 (41) was used to calculate differences in coverage for individual CAZymes, KEGG orthologs and RUGs. UpSet graphs were constructed using the UpSetR package (42). Taxonomies were assigned to paired sequence reads with Kraken (43) using a custom kraken database consisting of RefSeq complete genomes with our RUGs and the rumen superset (20) added. Prior to statistical analyses (excluding DeSeq2) and graph construction, data was subsampled. For RUGs, subsampling to the lowest sample coverage was performed. CAZymes and KEGG orthologs were subsampled to the lowest sample abundance.

## Results

### Construction of RUGs from rumen sequencing data

We produced 979G of Illumina sequencing data from 12 samples then performed a metagenomic assembly of single samples and a co-assembly of all samples. This created a set of 391 dereplicated genomes (99% ANI (average nucleotide identity)) with estimated completeness ≥80% and estimated contamination ≤10% (**Additional File 1**: **Fig 1**). 284 of these genomes were produced from the single-sample assemblies and 107 were produced from the co-assemblies. 172 genomes were >90% complete with contamination <5%, and would therefore be defined as high-quality draft genomes by Bower et al. (44). The distribution of these RUGs between our samples can be found in **Additional file 2** (based on coverage). **Additional file 3** contains the predicted taxonomic assignment for each RUG while **Fig 1** shows a phylogenetic tree of the genomes. The tree is dominated by the *Bacteroidota* (136 RUGs: All order *Bacteroidales*) and the *Firmicutes_A* (121 RUGs), followed by lesser numbers of the *Firmicutes_C* (40 RUGs), *Synergistota* (20 RUGs: All family *Aminobacteriaceae), Firmicutes* (19 RUGs), *Proteobacteria* (15 RUGs), *Cyanobacteriota* (9 RUGs: All family *Gastranaerophilaceae), Actinobacteriota* (7 RUGs), *Euryarchaeota* (7 RUGs: All family *Methanobacteriaceae), Spirochaetota* (5 RUGs), *Elusimicrobiota* (3 RUGs: All family *Endomicrobiaceae), UBP6* (3 RUGs: All genus *UBA1177), Fibrobacterota* (2 RUGs: All genus *Fibrobacter), Riflebacteria* (2 RUGs: All family *UBA8953), Chloroflexota* (1 RUGs: family *Anaerolineaceae*) and *Desulfobacterota* (1 RUGs: genus *Desulfovibrio*). All members of the phylum *Firmicutes_A* belonged to the *Clostridia* class: orders *4C28d-15* (n=9), *CAG-41* (n=3), *Christensenellales* (n=4), *Lachnospirales* (n=56), *Oscillospirales* (n=45), *Peptostreptococcales* (n=2) and *Saccharofermentanales* (n=2). *Firmicutes_C* contains the orders *Acidaminococcales* (n=8) and *Selenomonadales* (n=32). The phylum *Firmicutes* contained the orders *Acholeplasmatales* (n=3), *Erysipelotrichales* (n=1), *Izimaplasmatales* (n=1), *ML615J-28* (n=1), *Mycoplasmatales* (n=1). *RFN20* (n=7) and *RF39* (n=5), The *Actinobacteria* contained the orders *Actinomycetales* (n=1) and *Coriobacteriales* (n=6). The *Proteobacteria* phylum contains the orders *Enterobacterales* (n=4), *Paracaedibacterales* (n=1), *RF32* (n=8) and *UBA3830* (n=2). The *Spirochaetota* contains the orders *Sphaerochaetales* (n=1) and *Treponematales* (n=4).

**Fig 1:**
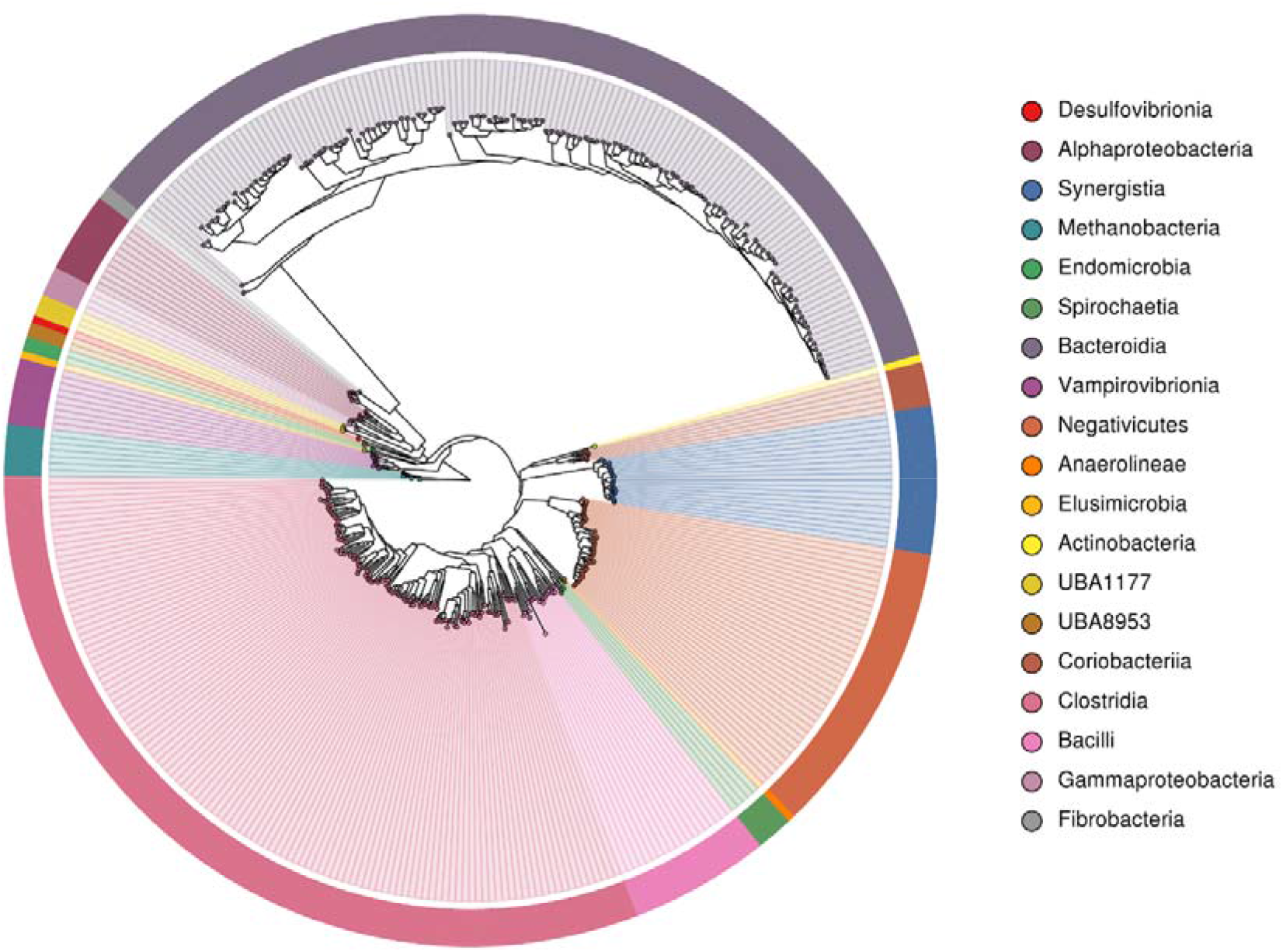
Phylogenetic tree of the 391 draft microbial genomes from rumen samples, labelled by taxonomic class. Taxonomies were defined by GTDB-Tk.

After sub-sampling, we found that samples from different ruminant species clustered significantly separately by abundance of RUGs (PERMANOVA: P = 3e-05). This may be due to the fact that the vast majority of RUGs were only found in a single host species (**Fig 2**), including 111 RUGs in red deer, 78 RUGs in reindeer, 40 RUGs in cow and 31 RUGs in sheep. Only 3 RUGs were found in ≥1X average coverage in all species: uncultured *Bacteroidaceae sp. RUG30019*, uncultured *Prevotella sp. RUG30028* and uncultured *Prevotella sp. RUG30114*.

We compared our RUGs to microbial genomes which had previously been sequenced from the rumen to determine if we had discovered any novel strains or species. We dereplicated our RUGs at 99% and 95% ANI to a “superset” of genomes containing rumen RUGs previously produced by our group (20), Hess et al. (11), Parks et al. (45), Solden et al. (46) and Svartström et al. (47) and the genomes from the Hungate collection (17). After dereplication at 99% and the removal of any RUGS with ≥99% ANI to an existing genome (as assigned by GTDB-Tk) or which clustered with members of the superset, 372 of our RUGs remained, representing putative novel strains. After dereplication at 95% and the removal of any RUGS with ≥95% ANI to an existing genome (assigned by GTDB-tk) or which clustered with members of the superset, 279 of our RUGs remained, representing putative novel species. The majority of these species originated from single-sample assemblies: 110 from red deer samples, 68 from reindeer samples, 23 from sheep samples and 1 from cattle samples, suggesting that many novel microbial species remain to be discovered from non-cattle ruminant hosts. These novel species are taxonomically diverse, with members belonging to the phyla *Bacteroidota* (n = 97), *Firmicutes_A* (n = 85), *Firmicutes_C* (n = 27), *Firmicutes* (n = 16), *Synergistota*(n = 14), *Proteobacteria* (n = 11), *Cyanobacteriota* (n = 9), *Actinobacteriota* (n = 5), *Spirochaetota* (n = 4), *Euryarchaeota* (n = 3), *Elusimicrobiota* (n = 3), *Riflebacteria* (n = 2), *Chloroflexota* (n = 1), *Desulfobacterota* (n = 1) and *UBP6* (n = 1).

31 of our total RUGs were able to be taxonomically identified to species level and these contain bacteria which are commonly isolated from the rumen including novel strains of *Bacteroidales bacterium UBA1184* (45), *Bacteroidales bacterium UBA3292* (45), *Butyrivibrio fibrisolvens, Escherichia coli, Fibrobacter sp. UWB2* (48), *Lachnospiraceae bacterium AC3007* (17), *Lachnospiraceae bacterium UBA2932* (45), *Methanobrevibacter sp. UBA188* (45), *Methanobrevibacter sp. UBA212* (45), *Prevotella sp. UBA2859* (45), *Ruminococcaceae bacterium UBA3812* (45), *Ruminococcus sp. UBA2836* (45), *Sarcina sp. DSM 11001* (17), *Selenomonas sp. AE3005* (17), *Succiniclasticum ruminis* and *Succinivibrio dextrinosolvens*.

### Comparing microbial taxonomies, CAZymes and KEGG orthologs between ruminant species

We assigned taxonomies to paired sequence reads using our custom kraken database containing RefSeq complete genomes, our RUGs, and the superset of rumen isolated microbial genomes. After subsampling we compared the abundance of members of the microbiota in different ruminant species at multiple taxonomic levels. Averaging reads across rumens species, the vast majority of reads mapped to bacteria (Sheep: 97%, Cow: 97%, Reindeer: 92%, Red deer: 98%) with smaller amounts of archaea (Sheep: 2.3%, Cow: 2.1%, Reindeer: 6.3%, Red deer: 1.9%) and Eukaryota (Sheep: 0.23%, Cow: 1.3%, Reindeer: 1.8%, Red deer: 0.56%). Eukaryota reads originated primarily from fungi and protists. In all ruminants, *Bacteroidetes* was the most abundant phylum (Sheep: 64%, Cow: 65% Reindeer: 54% Red deer: 52%), with *Firmicutes* being the second most abundant (Sheep: 29%, Cow: 26% Reindeer: 26% Red deer: 38%). Using PERMANOVA, significant differences in the abundance of taxonomies between ruminant species were found at both high (Kingdom: P = 0.01058, Phylum: P = 0.00017) and low (Family: P = 1e-05, Genus: P = 3e-05) taxonomic levels (**Additional File 1: Fig 2**).

**Fig 2:**
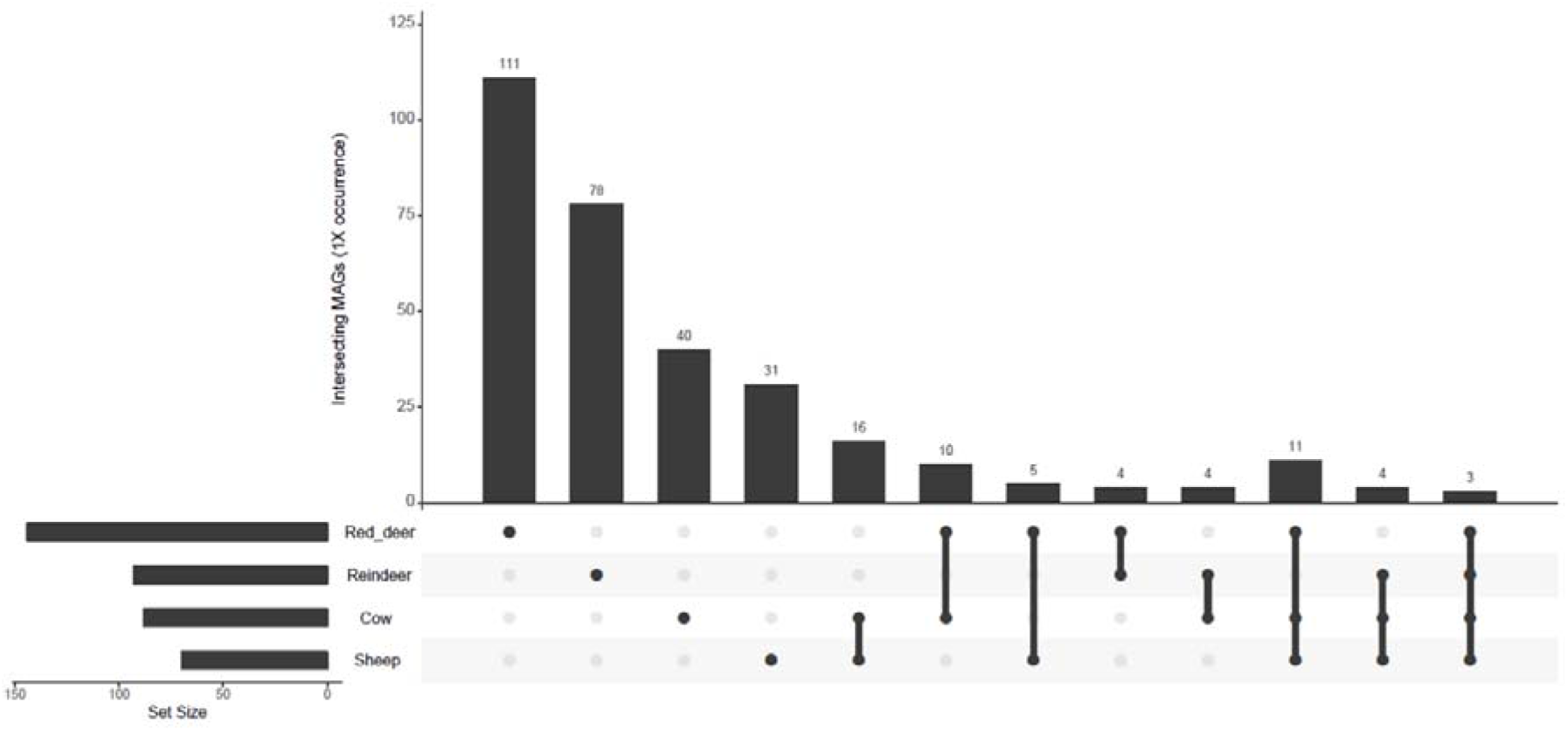
UpSetR graph showing the number of shared microbial genomes at average 1X coverage (after sub-sampling to equal depth) within four ruminant species.

We also compared the abundance of genes encoding for specific CAZymes between species. These enzymes are responsible for the synthesis, binding and metabolism of carbohydrates. The carbohydrate esterases (CEs), glycoside hydrolases (GHs), glycosyltransferases (GTs) and polysaccharide lyases (PLs) act to degrade cellulose, hemicellulose and other carbohydrates which could otherwise not be digested by the host. Non-catalytic carbohydrate-binding modules (CBMs) bind to specific carbohydrates, increasing the efficiency of enzymatic degradation (49). The auxiliary activities (AAs) redox enzymes are reclassified CBMs which are lytic polysaccharide monooxygenases (50). In our samples we found the following numbers of these CAZyme families: 6 AAs redox enzymes, 39 CBMs, 14 CEs, 191 GHs, 61 GTs and 27 PLs. The ten most abundant GHs in the different ruminant species were: for cows GH2, GH3, GH31, GH97, GH28, GH51, GH43_10, GH105, GH10 and GH95; for sheep GH2, GH3, GH28, GH31, GH97, GH32, GH51, GH77, GH78 and GH95; for red deer GH2, GH3, GH31, GH97, GH77, GH32, GH51, GH109, GH28 and GH78; and for reindeer GH2, GH3, GH92, GH109, GH97, GH13, GH31, GH78, GH28 and GH77. Different ruminant species were found to have significantly differently abundant CAZyme genes (PERMANOVA: P = 1e-05, Additional File 1: Fig 3). However, it should be noted that the vast majority of CAZyme families were found in all sample types (Fig 3), indicating that there exists a set of CAZymes which are present across ruminant species consuming different diets and living in vastly different conditions.

**Fig 3:**
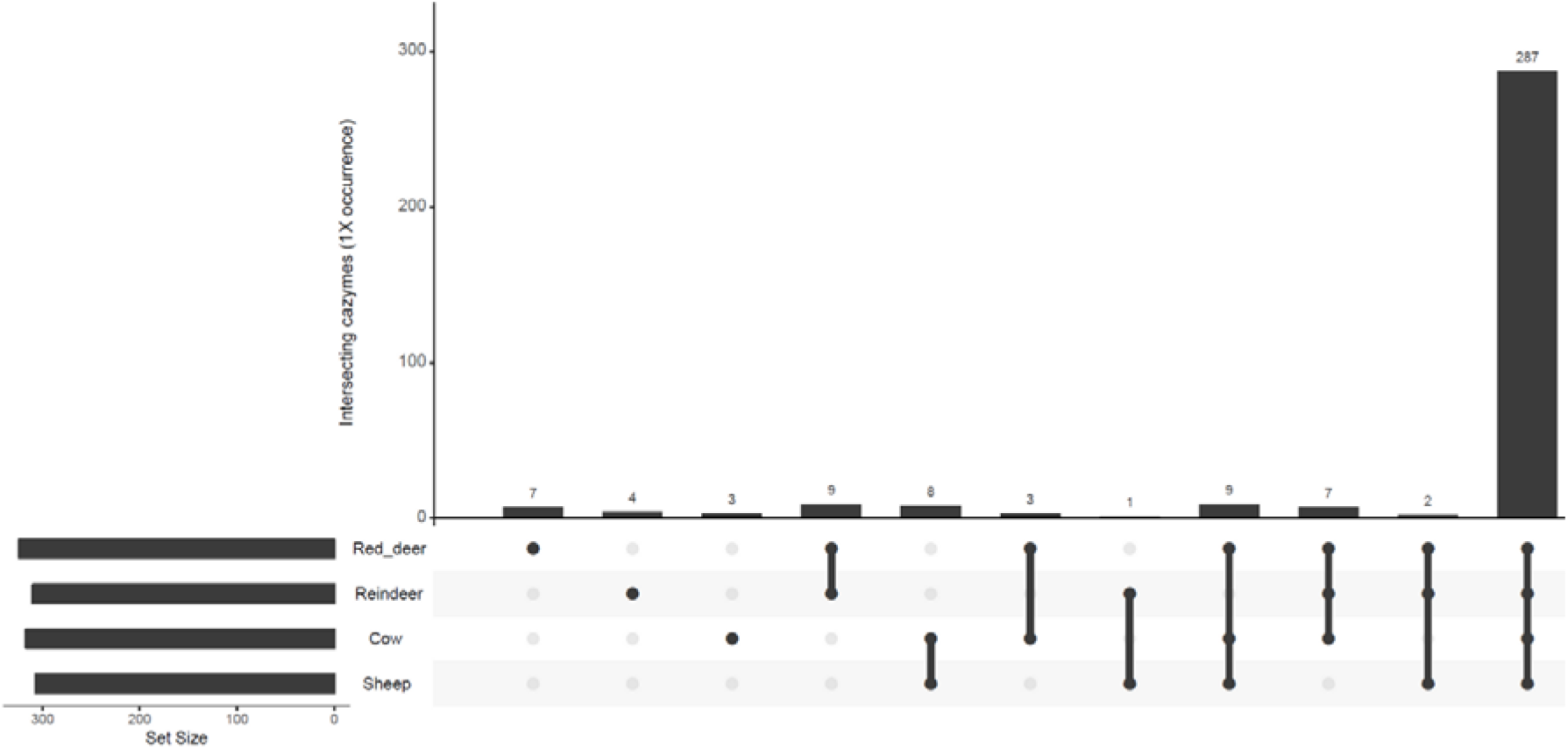
UpSetR graph showing the number of shared CAZyme families at average 1X coverage within four ruminant species.

DeSeq2 was used to identify specific CAZymes which were significantly more abundant in one ruminant species vs another (**Additional file 4**). Those CAZymes which were consistently more abundant in specific species when compared to other species are listed in **Tables 1–4.**

**Table 1:** CAZymes (Carbohydrate-active enzymes) which were consistently more abundant in cattle.

| CAZyme | Deer -<br>adjusted p-value | Sheep -<br>adjusted p-value | Reindeer -<br>adjusted p-value |
| --- | --- | --- | --- |
| Carbohydrate-binding modules |  |  |  |
| CBM11 | 0.000113 | 1.81E-05 | 8.41E-05 |
| CBM22 | 0.000996 | 0.000958 | 0.004915 |
| CBM25 | 0.000571 | 0.010859 | 0.005573 |
| CBM6 | 4.32E-11 | 0.02129 | 1.91E-25 |
| CBM74 | 4.51E-06 | 0.009696 | 0.00389 |
| Carbohydrate esterases |  |  |  |
| CE15 | 1.03E-07 | 0.000403 | 0.000357 |
| CE6 | 6.57E-13 | 0.000634 | 1.07E-27 |
| Glycoside hydrolases |  |  |  |
| GH105 | 8.55E-10 | 0.0195 | 1.22E-14 |
| GH11 | 0.000134 | 0.000687 | 2.24E-05 |
| GH115 | 0.000737 | 0.018632 | 1.91E-17 |
| GH13_28 | 7.54E-06 | 0.012113 | 0.042438 |
| GH130 | 1.02E-08 | 2.69E-06 | 0.02559 |
| GH146 | 0.011892 | 0.032424 | 3.57E-06 |
| GH31 | 4.20E-08 | 0.000243 | 7.31E-10 |
| GH35 | 3.11E-06 | 0.003975 | 5.19E-10 |
| GH36 | 0.001473 | 0.002527 | 1.22E-13 |
| GH43_1 | 5.48E-12 | 6.03E-05 | 2.36E-22 |
| GH43_24 | 3.66E-06 | 0.000849 | 0.022236 |
| GH43_29 | 5.69E-10 | 0.000779 | 3.84E-65 |
| GH43_35 | 1.76E-06 | 0.018491 | 4.78E-29 |
| GH43_5 | 4.79E-16 | 0.011903 | 2.98E-40 |
| GH43_7 | 2.36E-09 | 0.012453 | 1.60E-36 |
| GH45 | 1.13E-05 | 0.000541 | 0.001412 |
| GH5_10 | 9.78E-05 | 0.000118 | 0.000164 |
| GH5_38 | 1.36E-11 | 0.022559 | 1.78E-37 |
| GH5_39 | 6.13E-05 | 0.000849 | 0.02559 |
| GH5_52 | 0.00487 | 0.015377 | 0.003089 |
| GH51 | 6.44E-08 | 0.011587 | 2.36E-12 |
| GH67 | 1.50E-11 | 0.001929 | 1.45E-31 |
| GH8 | 4.01E-10 | 0.000212 | 2.96E-31 |
| GH9 | 1.12E-06 | 1.08E-07 | 0.002527 |
| GH94 | 1.45E-05 | 5.96E-13 | 0.000853 |

**Table 2:** CAZymes (Carbohydrate-active enzymes) which were consistently more abundant in sheep.

| CAZyme | Cow - adjusted<br>p-value | Deer -<br>adjusted p-value | Reindeer -<br>adjusted p-value |
| --- | --- | --- | --- |
| Carbohydrate-binding modules |  |  |  |
| CBM27 | 0.019613 | 1.62E-07 | 1.13E-05 |
| Carbohydrate esterases |  |  |  |
| CE8 | 0.021156 | 1.65E-08 | 4.29E-14 |
| Glycoside hydrolases |  |  |  |
| GH144 | 6.75E-05 | 0.003343 | 0.034267 |
| GH30_8 | 0.026986 | 0.0002685 | 2.49E-32 |
| GH53 | 0.000983 | 6.98E-14 | 9.02E-14 |
| Glycosyltransferases |  |  |  |
| GT69 | 7.60E-05 | 5.21E-05 | 4.40E-06 |
| GT7 | 3.50E-08 | 0.0002904 | 0.000314 |
| Polysaccharide lyases |  |  |  |
| PL10 | 0.022559 | 2.52E-07 | 3.35E-22 |

**Table 3:** CAZymes (Carbohydrate-active enzymes) which were consistently more abundant in red deer.

| CAZyme | Cow - adjusted p-value | Reindeer - adjusted p-value | Sheep - adjusted p-value |
| --- | --- | --- | --- |
| Carbohydrate-binding modules |  |  |  |
| CBM34 | 2.18E-22 | 3.69E-13 | 0.049585 |
| CBM42 | 0.005095 | 0.003958 | 0.000579 |
| CBM54 | 2.87E-14 | 4.60E-07 | 0.000154 |
| CBM58 | 5.65E-05 | 0.00122 | 0.000348 |
| Carbohydrate esterases |  |  |  |
| CE13 | 0.002535 | 5.30E-06 | 0.006324 |
| Glycoside hydrolases |  |  |  |
| GH13_20 | 3.56E-09 | 0.014494 | 0.021084 |
| GH13_29 | 2.22E-40 | 4.90E-08 | 0.006428 |
| GH13_4 | 9.39E-08 | 0.000614 | 0.000579 |
| GH147 | 0.00095 | 0.000495 | 0.000842 |
| GH148 | 5.24E-10 | 1.92E-08 | 0.0105 |
| GH24 | 5.09E-23 | 0.00023 | 1.24E-05 |
| GH43 | 0.010658 | 0.000118 | 0.024375 |
| GH43_11 | 4.01E-10 | 0.004229 | 0.024919 |
| GH43_21 | 3.74E-06 | 0.004636 | 0.000375 |
| GH5_44 | 8.22E-23 | 0.000382 | 0.001021 |
| Glycosyltransferases |  |  |  |
| GT23 | 2.31E-11 | 0.00713 | 0.036234 |
| Polysaccharide lyases |  |  |  |
| PL4 | 4.20E-69 | 1.94E-19 | 2.12E-44 |

**Table 4:** CAZymes (Carbohydrate-active enzymes) which were consistently more abundant in reindeer.

| **CAZyme** | **Cow - adjusted p-value** | **Deer - adjusted p-value** | **Sheep - adjusted p-value** |
| --- | --- | --- | --- |
| Carbohydrate-binding modules |  |  |  |
| CBM32 | 4.15E-29 | 8.21E-08 | 4.63E-11 |
| CBM41 | 0.00392 | 1.73E-05 | 1.71E-11 |
| CBM62 | 5.74E-06 | 1.52E-05 | 1.95E-09 |
| CBM66 | 2.12E-29 | 1.66E-13 | 3.43E-11 |
| CBM67 | 6.33E-11 | 5.31E-05 | 0.000125 |
| CBM68 | 5.01E-05 | 8.99E-06 | 0.000182 |
| CBM9 | 8.88E-13 | 1.44E-11 | 2.22E-12 |
| Carbohydrate esterases |  |  |  |
| CE3 | 1.68E-10 | 7.47E-05 | 0.00062 |
| CE9 | 5.83E-19 | 7.37E-05 | 0.007305 |
| Glycoside hydrolases |  |  |  |
| GH109 | 1.80E-18 | 9.23E-08 | 1.92E-13 |
| GH117 | 1.25E-05 | 0.004221 | 0.003013 |
| GH123 | 0.003063 | 0.000606 | 0.001734 |
| GH125 | 4.14E-19 | 1.33E-16 | 1.59E-15 |
| GH128 | 2.16E-14 | 9.97E-07 | 4.18E-05 |
| GH13 | 0.004306 | 1.47E-07 | 0.007748 |
| GH13_16 | 7.00E-05 | 0.00063 | 3.51E-05 |
| GH13_2 | 0.005224 | 0.0006 | 0.026906 |
| GH13_36 | 0.002131 | 0.003061 | 0.027897 |
| GH13_40 | 0.000141 | 1.57E-05 | 4.25E-07 |
| GH133 | 0.030687 | 1.47E-07 | 0.000337 |
| GH16 | 7.62E-10 | 6.74E-06 | 0.000368 |
| GH18 | 1.28E-07 | 2.39E-07 | 0.000111 |
| GH20 | 6.60E-12 | 0.000614 | 1.03E-05 |
| GH38 | 9.43E-46 | 1.51E-18 | 3.04E-43 |
| GH39 | 2.69E-07 | 0.000556 | 1.43E-10 |
| GH43_26 | 1.57E-05 | 0.000556 | 0.000556 |
| GH43_28 | 1.35E-16 | 3.00E-08 | 4.40E-06 |
| GH43_3 | 5.69E-08 | 7.45E-07 | 8.84E-19 |
| GH43_31 | 5.43E-07 | 0.000583 | 2.26E-07 |
| GH43_33 | 1.58E-06 | 2.24E-05 | 1.01E-13 |
| GH5 | 1.57E-22 | 1.80E-08 | 1.55E-15 |
| GH5_35 | 8.84E-13 | 0.001397 | 4.49E-06 |
| GH57 | 0.003543 | 0.005297 | 9.37E-05 |
| GH59 | 0.000148 | 0.003958 | 0.008335 |
| GH64 | 1.91E-09 | 1.87E-07 | 2.57E-09 |
| GH76 | 2.09E-20 | 1.55E-27 | 3.29E-16 |
| GH85 | 0.001061 | 0.002853 | 0.008951 |
| GH87 | 4.47E-10 | 2.62E-07 | 1.42E-07 |
| GH88 | 0.001577 | 0.00356 | 0.006143 |
| GH92 | 8.02E-20 | 3.58E-16 | 8.82E-23 |
| GH93 | 1.65E-08 | 4.49E-06 | 3.88E-11 |
| GH99 | 2.06E-07 | 2.73E-09 | 0.000163 |
| Glycosyltransferases |  |  |  |
| GT1 | 1.86E-05 | 1.86E-10 | 1.93E-05 |
| GT10 | 2.68E-08 | 0.012407 | 0.005445 |
| GT2_Glyco_tranf_2_4 | 5.94E-21 | 0.000115 | 2.57E-07 |
| GT3 | 0.008785 | 0.001394 | 0.001478 |
| GT39 | 0.002494 | 0.021868 | 0.003876 |
| GT4 | 7.61E-09 | 0.000752 | 0.018262 |
| GT47 | 0.000453 | 0.011114 | 0.002988 |
| GT66 | 2.68E-06 | 5.25E-05 | 0.008382 |
| GT70 | 9.38E-07 | 0.000433 | 0.001094 |
| GT77 | 0.002527 | 0.014307 | 0.009975 |
| GT8 | 5.78E-21 | 0.000151 | 0.02682 |
| GT81 | 1.55E-06 | 8.26E-06 | 0.025595 |
| GT83 | 4.25E-39 | 4.13E-05 | 1.77E-06 |
| Polysaccharide lyases |  |  |  |
| PL17_2 | 2.55E-05 | 0.000285 | 0.00041 |
| PL22 | 1.27E-06 | 0.005756 | 0.000133 |
| PL8 | 0.003063 | 0.002491 | 0.006191 |

CAZymes are often found organised into Polysaccharide Utilization Loci (PUL) which comprise a set of genes that enable the binding and degradation of specific carbohydrates or multiple carbohydrates. We used the software PULpy to predict PULs which were present in our *Bacteroidales* RUGs. Of the 136 RUGs which belong to the taxonomy *Bacteroidales*, 112 contain putative PULs. Within these RUGs we identified 970 PULs, with numbers of PULs per RUG ranging from 1 to 35. The largest quantity of PULs originating from one RUG was 35 from uncultured *Bacteroidales sp. RUG30227*; these encoded a wide range of CAZymes. This RUG was more abundant in reindeer samples than samples from other ruminants. Of the 970 PULs, 332 of these were a single susC/D pair. A summary of identified PULs can be found in **Additional file 5 (Additional File 1: Fig 4**).

**Fig 4:**
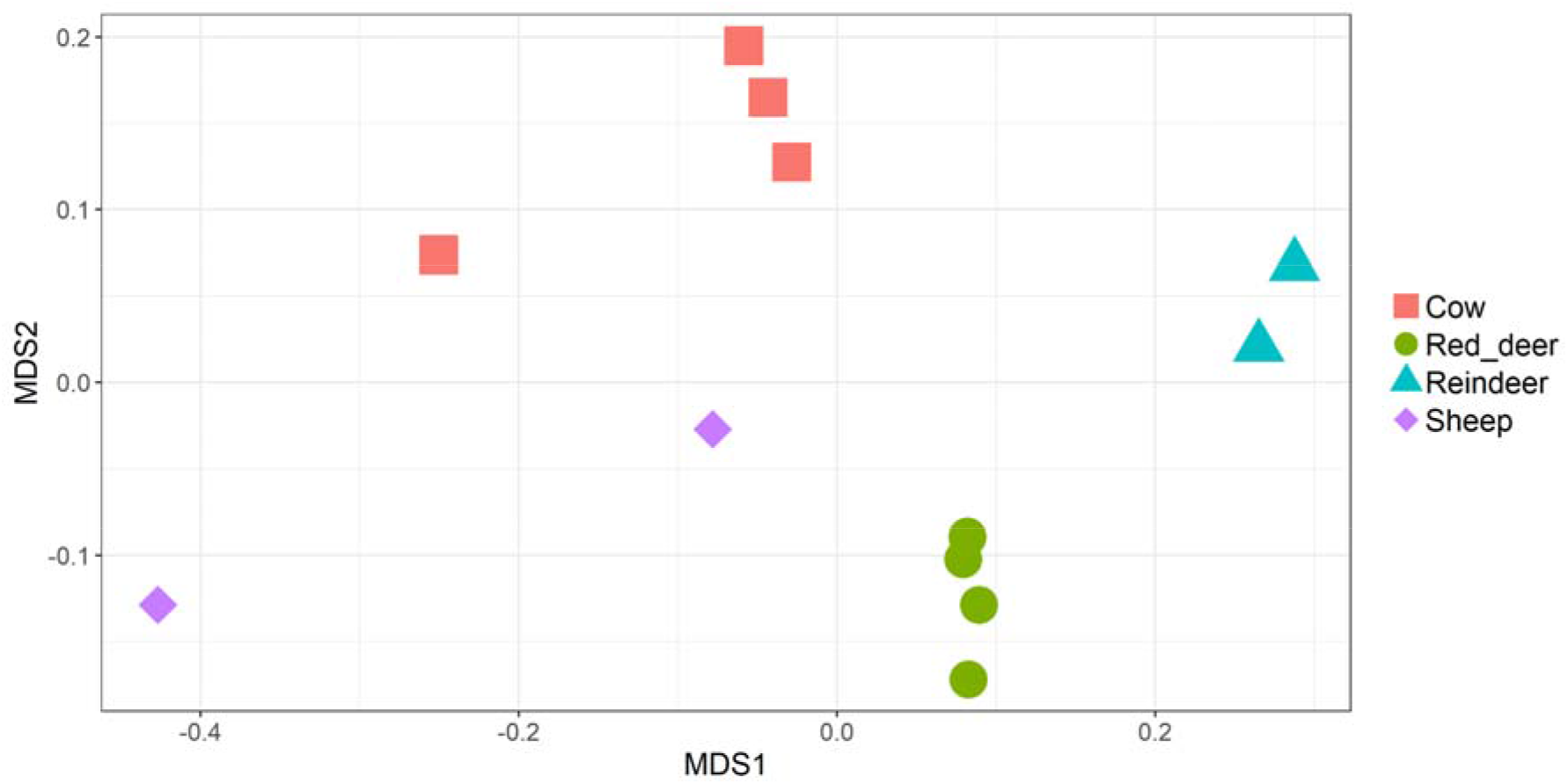
NMDS of ruminal samples clustered by abundance of KEGG orthologs, using Bray–Curtis dissimilarity values (PERMANOVA; P = 1e-05).

We also examined the abundance of genes which belonged to specific KEGG orthologs. KEGG orthologs represent a wide range of molecular functions and are defined by a network-based classification. We found that, as for CAZymes, ruminant species clustered significantly by the abundance of genes with specific KEGG orthologs (PERMANOVA: P = 1e-05, **Fig 4**) and that the vast majority of orthologs were found in all ruminant species (**Fig 5**). However, the large amount of orthologs (n=729) which were only found in the two domesticated species (cattle and sheep) is also worthy of note. It should also be noted that the two sheep samples did not cluster visually to the same extent as the samples originating from the other ruminant speices (**Fig 4**). DeSeq2 was used to identify many KEGG orthologs which were significantly more abundant in one ruminant species vs another (**Additional file 6**). Those orthologs which were consistently more abundant in specific ruminant species (Adjusted p-value <0.05) are listed in **Additional file 7.**

**Fig 5:**
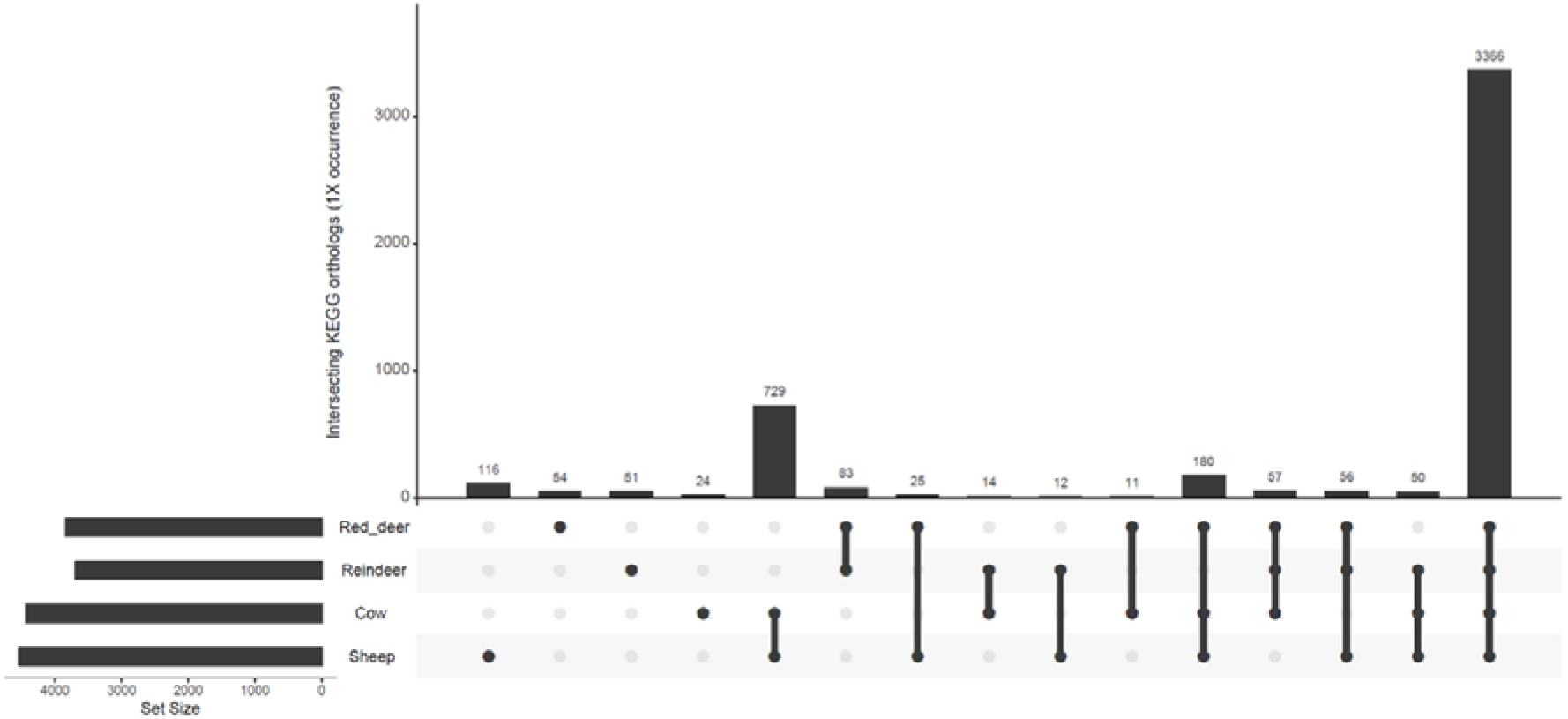
UpSetR graph showing the number of shared KEGG orthologs families at average 1X coverage within four ruminant species.

## Discussion

The rumen microbiota plays a crucial role in the ability of ruminants to efficiently digest feed while the rumen microbiota and their products also have a potential use in diverse industrial applications. The ruminal microbiota of red deer and reindeer have previously been studied using 16S rRNA gene sequencing (51–54). However, metagenomic studies of these species are limited, with only one study in reindeer (55) and no studies in red deer.

In this study we constructed 391 rumen microbial genomes from metagenomic data from cattle, sheep, red deer and reindeer. We assigned taxonomies to our RUGs using GTDB-Tk rather than NCBI based taxonomies as this improves the classification of uncultured bacteria due to the use of a genome-based taxonomy (32). We have also previously found less need to manually correct taxonomic assignments when using GTDB-Tk (21). Our microbes predominantly belonged to the *Bacteroidota* and *Firmicutes_A*, with lesser numbers of 14 other phyla. We dereplicated our genomes alongside a superset of rumen bacterial genomes (20) and used the results output by GTDB-Tk to identify RUGs which represent novel microbial strains and species. Amongst our genomes we identified 372 novel strains and 279 novel species. These microbes were taxonomically diverse, belonging to 15 phyla. Only 31 RUGs were assigned an identity at species level.

The vast majority of our total RUGs were only present on average at ≥1 coverage in one ruminant species. However, we found that at higher taxonomic levels taxonomies were shared between sample types. When comparing the abundance of taxonomies between samples we found that ruminant species clustered separately by both higher (kingdom and phylum) and lower (family and genus) taxonomic levels. We are aware that the sample sizes for our study are small and therefore any conclusions about differences between the microbiota of ruminant species should be drawn cautiously. However, our data are supportive of the hypothesis that there are host species-specific rumen microbiota at the strain and species level but that these differences do not necessarily translate into large differences in the types of CAZymes expressed.

While we found that there were significant differences between the abundances of CAZymes between ruminant species, most CAZymes were present in all ruminant species. These results also reflected those which we found when analysing the abundance of KEGG orthologs. We also identified 970 PULs in our *Bacteriodales* RUGs, with numbers of PULs per genome ranging from 1 to 35. The RUG containing 35 PULs was found most abundantly in reindeer samples, emphasising the potential for the discovery of novel carbohydrate-active enzymes in lesser studied ruminant species, as also highlighted by a previous study which identified multiple PULs in metagenomic samples from reindeer (55). Unfortunately due to the nature of our samples, with red deer and reindeer samples originating from animals eating a non-regimented diet, we are not able to provide metadata as to the exact nutritional composition of our animals’ diets, therefore a more in depth analysis of dietary carbohydrates vs CAZyme/PUL abundance is not possible.

While several thousand RUGs have previously been published that originate from the rumen microbiota, the vast majority of these originate from cattle. By investing more effort in exploring the metagenome of less well studied ruminants we will be able to identify even more microbes and microbial products that are of industrial-interest. In conclusion, we present a dataset of RUGs from four ruminant species which can be used as a reference dataset in future metagenomic studies and to aid in the design of culture based studies.

## Author statements

### Authors and contributors

LG contributed to methodology, formal analysis, data curation, writing (original draft preparation) and visualisation. BG and RJW contributed to conceptualisation, methodology, investigation, resources and writing (review and editing). RJW also contributed to supervision and project administration. MW contributed to conceptualisation, methodology, formal analysis, writing (review and editing), visualisation, supervision and project administration. All authors read and approved the final manuscript.

### Ethical approval

Cattle projects were carried out under Home Office PPL 30/2579. Sheep experimentation was carried out under the conditions set out by UK Home Office licence no. 604028, procedure reference number 8.

### Consent for publication

Not applicable.

## Acknowledgments

BG thanks the Higher Education Council of the Turkish Republic for the award of a travelling fellowship. We thank Bob Mayes and Dave Hamilton of the James Hutton Institute for their permission and help in sampling the sheep digesta, Kevin Shingfield for the provision of reindeer digesta samples, and Euan Munro for red deer digesta samples. We also thank Nest McKain of RI for technical assistance.

## Supporting data

**Additional file 1.docx: S1 Figures** - Supplementary figures 1 to 4.

**Additional file 2.xls: Dataset 1** - Average coverage of RUGs in all samples. Coverage was calculated by mapping the RUG scaffolds to adaptor-trimmed Illumina reads.

**Additional file 3.xls: Dataset 2** - Description of each RUG, including taxonomy, novelty, genome completeness, genome contamination, genome size, number of ambiguous bases, number of scaffolds, number of contigs, N50 (scaffolds), N50 (contigs), mean scaffold length, mean contig length, longest contig and GC content.

**Additional file 4.xls: Dataset 3** - CAZymes which were more abundant in sheep, cattle, reindeer or red deer (DeSeq2: Adjusted p-value <0.05).

**Additional file 5: Dataset 4** - Summary of polysaccharide utilisation loci found in *Bacteroidales* RUGs.

**Additional file 6: Dataset 5** - KEGG orthologs which were more abundant in sheep, cattle, reindeer or red deer (DeSeq2: Adjusted p-value <0.05).

**Additional file 7: Dataset 6** - KEGG orthologs which were consistently more abundant in sheep, cattle, reindeer or red deer (DeSeq2: Adjusted p-value <0.05).

