## Additional file 1 for "Metagenomic analysis of the cow, sheep, reindeer and red deer rumen"

**Supplementary figure 1:**

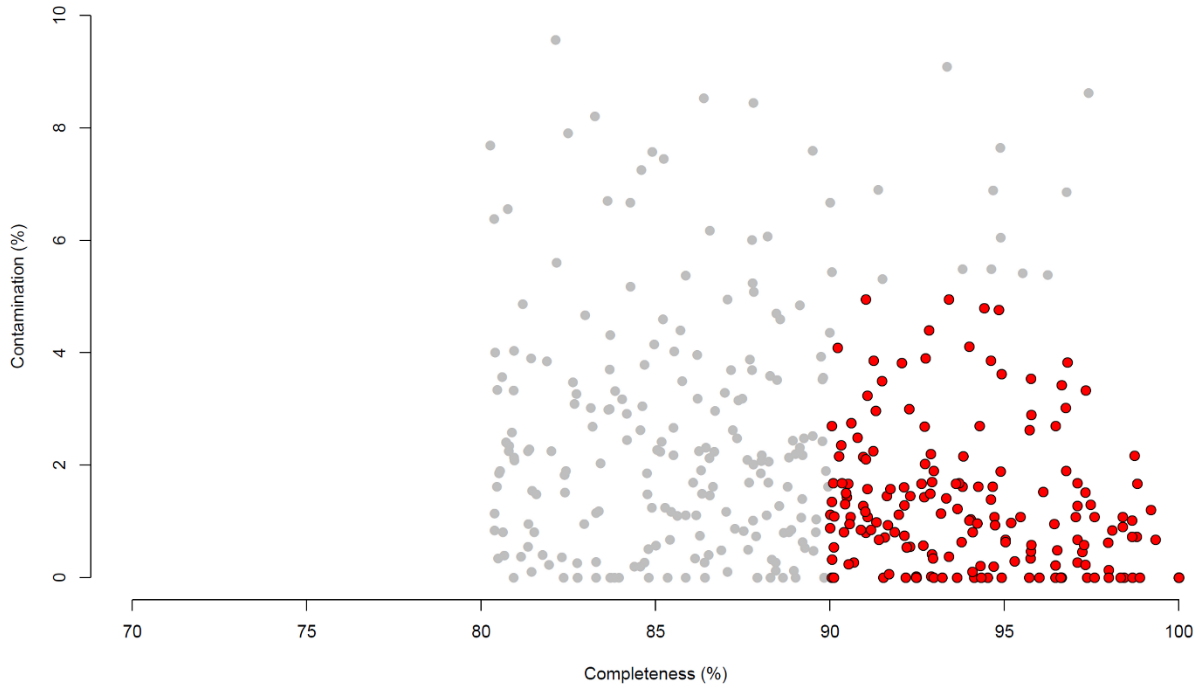


**Contamination and completeness (defined by CheckM software) of 391 dereplicated metagenome-assembled-genomes from rumen samples. Grey: genomes which are 80-90% complete with 5-10% contamination. Red: genomes which are >90% complete with <5% contamination.**

**Supplementary figure 2:**

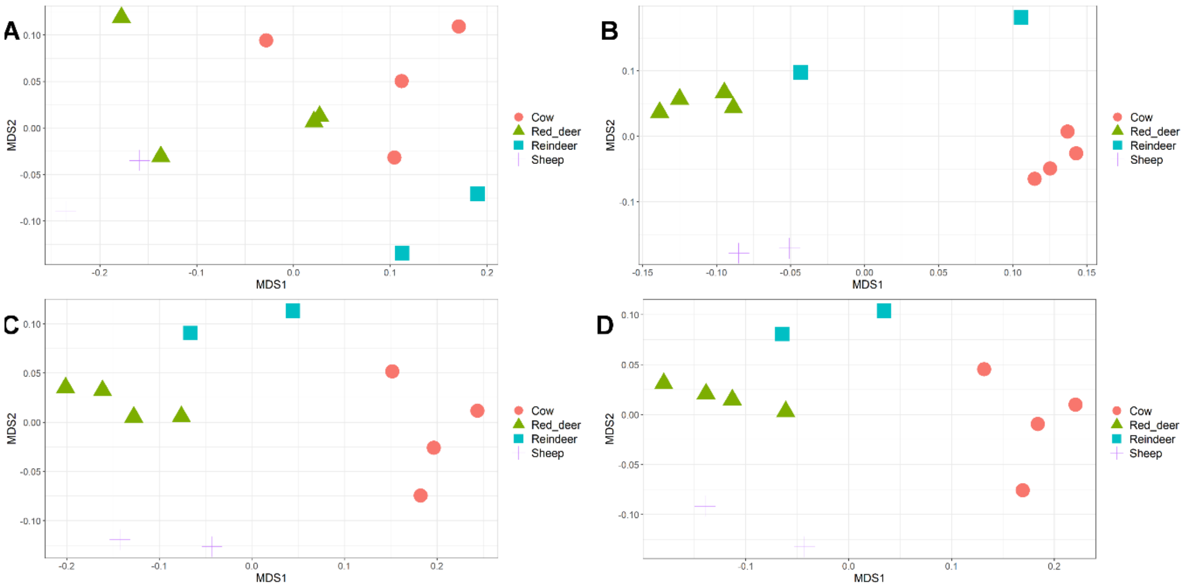


**NMDS of ruminal samples clustered by abundance of taxonomies, using Bray–Curtis dissimilarity values. A: Kingdom (PERMANOVA; P =0.01058), B: Phylum (PERMANOVA; P= 0.00017), C: Family (PERMANOVA; P =1e-05), D: Genus (PERMANOVA; P =3e-05).**

**Supplementary figure 3:**

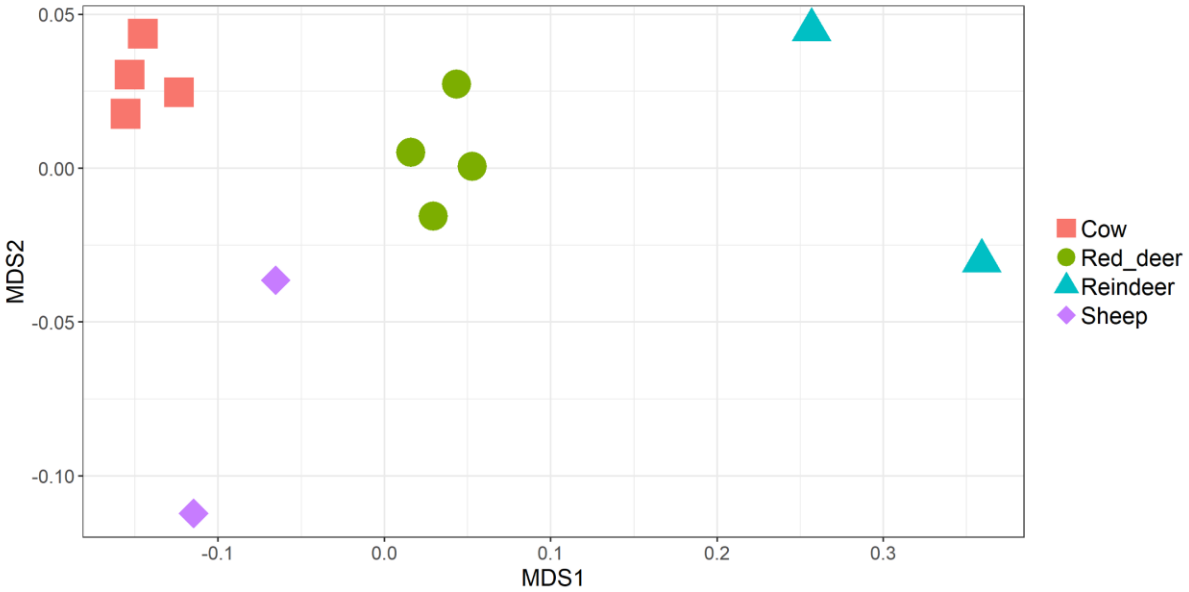


**NMDS of ruminal samples clustered by abundance of CAZymes, using Bray–Curtis dissimilarity values (PERMANOVA; P = 1e-05).**

**Supplementary figure 4**

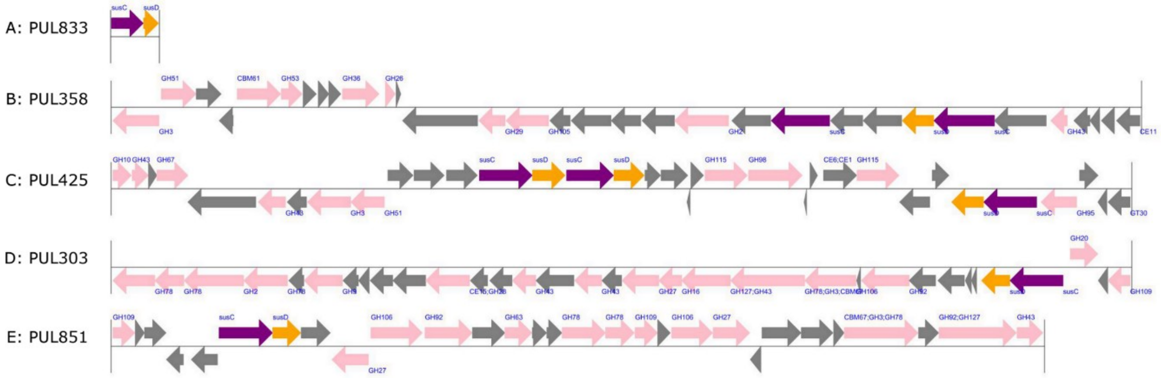


**Several predicted polysaccharide utilisation loci (PULs). A: The most common PUL consisting of only a susC/D pair. B-E: The 4 largest PULs identified in our dataset.**
